# TLR2 on CD4+ and CD8+ T cells promotes late control of *Mycobacterium tuberculosis* infection

**DOI:** 10.1101/2021.05.19.444905

**Authors:** Scott M. Reba, Qing Li, Sophia Onwuzulike, Nancy Nagy, Kyle Parker, Katharine Umphred-Wilson, Supriya Shukla, Clifford V. Harding, W. Henry Boom, Roxana E. Rojas

**Author notes:** Shared senior authors (Clifford V. Harding, W. Henry Boom, Roxana E. Rojas).

## Abstract

Although a role for TLR2 on T cells has been indicated in prior studies, *in vivo* stimulation of TLR2 on T cells by *Mtb* and its impact on *Mtb* infection has not been tested. Furthermore, it is not known if the enhanced susceptibility to *Mtb* of *Tlr2* gene knockout (ko) mice is due to its role in macrophages, on T cells or both. To address TLR2 on T cells, we generated *Tlr2*^*fl/fl*^*xCd4*^*cre/cre*^ mice, which lack expression of TLR2 on both CD4 and CD8 T cells, to study the *in vivo* role of TLR2 on T cells after aerosol infection with virulent *Mtb*. Deletion of TLR2 in CD4+ and CD8+ T cells reduces their ability to be co-stimulated by TLR2 ligands for cytokine production. These include both pro-(IFN-γ, TNF-α) and anti-inflammatory cytokines (IL-10). Deletion of TLR2 in T cells did not affect early control but did result in decreased late control of *Mtb* in the lungs of infected mice. This suggests that T cell co-stimulation by mycobacterial TLR2 ligands *in vivo* is important for control of infection during the chronic phase of *Mtb* infection in the lung.

## Introduction

*Mycobacterium tuberculosis* (*Mtb*) is one of the most successful pathogens, infecting one-third of the world’s population and is a major cause of adult mortality [1]. Classical MHC-restricted T cells have a central role in controlling *Mtb* infection (reviewed in [2]). T-cell activation requires two signals, signal 1 elicited by antigen peptide-MHC engagement of the TCR-CD3 complex, and signal 2 triggered when costimulatory receptor CD28 binds CD80/CD86 on antigen presenting cells (APC) (reviewed in [3]). Co-stimulation is required for naive T-cell priming, and important for optimal effector and memory T cell responses.

Besides constitutively expressed CD28, additional costimulatory receptors such as toll-like receptor 2 (TLR2) are induced after T cell activation [4]. TLRs are unique T cell costimulators because they are directly engaged by microbial ligands. *Mtb* is particularly enriched for TLR2 ligands [5], and TLR2 on human CD4+ T cells is expressed *in vivo* and interacts with *Mtb* lipoproteins and glycolipids [6, 7]. In combination with TCR-CD3 triggering, TLR2 engagement induces CD4+ T-cell proliferation and secretion of IL-2 and IFN-γ [6]. Engagement of TLR2 on CD4+ T-cells *in vivo* increases Th1-type cytokine secretion and proliferation [8]. Adoptively transferred *Mtb*-specific transgenic CD4+ T cells activated through TCR-CD3 plus TLR2 also were more effective in controlling *Mtb in vivo* than T cells activated via TCR-CD3 alone [8]. TLR2 also can serve as a co-stimulator for CD8+ T cells and enhance antiviral responses [9]. Its role in CD8 responses to *Mtb* is not known.

*Tlr2* gene knockout (ko) mice have reduced acquired immunity and increased bacterial growth after low-dose *Mtb* aerosol infection and increased mortality after high-dose infection [10, 11]. TLR2’s role in protecting the lung against *Mtb* could be due to its role in macrophages and APCs, T cells, other cell types, or a combination of these cells. Here we further determine the contribution of TLR2 expression on CD4+ and CD8+ T cells to control *Mtb* infection *in vivo*, using a T cell-specific *Tlr2*-deletion mouse model. We provide evidence that lack of TLR2 expression on T cells results in decreased control of late *Mtb* infection in the lung.

## Materials and Methods

### Mice

8-12 wk old female, C57BL/6J mice (WT) were purchased from Charles River Laboratories (North Wilmington, MA). CD4-Cre transgenic mice were purchased from Jackson Laboratories (#017336; C57Bl/6 background). Floxed TLR2 (*Tlr2*^fl/null^) C57Bl/6 mice were generated by Taconic (CWRU0002 Tlr2) by placing LoxP sites around exon 3 of *Tlr2* and bred for 8 generations to homozygous *Tlr2*^fl/fl^. CD4-Cre and *Tlr2*^fl/fl^ mice were crossed until homozygous *Tlr2*^*fl/fl*^*xCd4*^*cre/cre*^ mice were generated and then bred for 4 generations before use in experiments. Mice were housed under specific-pathogen-free conditions in ventilated microisolator cages (Lab Products, Inc., Maywood, NJ). The Institutional Animal Care and Use Committee at Case Western Reserve University approved all studies.

### Genetic Screening for presence and deletion of Tlr2

Genetic changes in transgenic mice were confirmed by gel electrophoresis. Tissue DNA was extracted with ReliaPrep gDNA Tissue Miniprep System (Promega) following. DNA was amplified with forward and reverse primers with Excellastart Dye Track Master Mix (Genessee) and analyzed by 4% polyacrylamide gel. The following primers were used in the generation of *Tlr2*^*fl/fl*^ mice and for assessing deletion of *Tlr2* in *Tlr2*^*fl/fl*^*xCd4*^*cre/cre*^ mice: for the 5’ *LoxP* site, *Tlr2*-flox forward, GCTTTCCGAGAACTTAACTCAGC (P1 in Fig 1A) and *Tlr2*-flox reverse, TGAACTAGATGTGGCCTATGGC (P2 in Fig. 1A). The 3’ *LoxP* was linked with *hGHpA31* and primer CCCAAGGCACACAAAAAACC used to detect exon 3 deletion (P3 in Fig. 1A). *Cre* forward, GCGGTCTGGCAGTAAAAACTATC and *cre* reverse, GTGAAACAGCATTGCTGTCACTT were used to detect CD4^cre/cre^ (https://www.jax.org/Protocol?stockNumber=017336&protocolID=27408). Primer pairs Mm00442346_m1 and Mm00607939_s1 from Invitrogen were used to measure TLR2 and actin-B mRNAs respectively.

**Figure 1.**
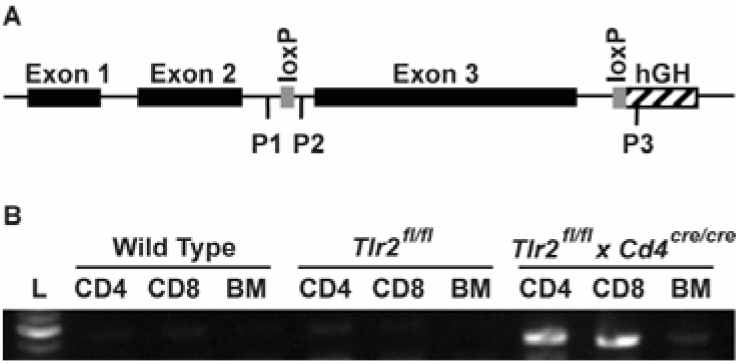
*Deletion of exon 3 of Tlr2 in CD4+ and CD8+ T cells in Tlr2*^*fl/fl*^*xCd4*^*cre/cre*^ *mice*. **A**. Location of LoxP sites around exon 3 of *Tlr2* in *Tlr2*^*fl/fl*^ mice, and location of primers (P1-3) used to confirm homozygosity of floxed *Tlr2* in *Tlr2*^*fl/fl*^ mice and exon 3 deletion in *Tlr2*^*fl/fl*^*xCd4*^*cre/cre*^ mice. **B**. Confirmation of deletion of exon 3 of Tlr2 in CD4+ and CD8+ T cells from *Tlr2*^*fl/fl*^*xCd4*^*cre/cre*^ mice. Upon deletion of exon 3, DNA PCR with primers P1 and P3 reveals a band in CD4+ and CD8+ T cells not present in CD4+ and CD8+ T cells of WT and *Tlr2*^*fl/fl*^ mice. Bone Marrow Derived Macrophages (BM in Figure 1b) from *Tlr2*^*fl/fl*^*xCd4*^*cre/cre*^ mice also do not reveal this band, confirming deletion of exon 3 of *Tlr2* only in T cells.

### TLR ligands and M. tuberculosis bacteria

Pam3Cys (EMC Micro-collections GmbH, Germany) was used as TLR2/1 agonist [14]. *Mtb* H37Rv (Colorado State University) was grown on 7H10 agar plates. Colonies were transferred to liquid GAS Media and grown to an absorbance of 1.1 (595 nm). Aliquots prepared in media supplemented with 20% glycerol were frozen at −80°C until use. Stock bacterial counts were determined by colony-forming unit (CFU) method.

### Cell culture

Purified resting CD4^+^ and CD8^+^ T cells were isolated from spleens with a one h adherence step, followed by negative selection using immune-magnetic cell sorting (Miltenyi, MACS). Purity was confirmed by flow cytometry. All *in vitro* experiments were performed with purified CD4^+^ and CD8+ T cells cultured in complete media, consisting of DMEM (BioWhittaker, Walkersville, MD.) supplemented with fetal calf serum (10% v/v, Gemini-Bioproducts, West Sacramento, CA; 100–106), penicillin (100 IU/mL), streptomycin (100 μg/mL), HEPES (100 mM), nonessential amino acids (100 mM), L-glutamine (2 mM), and 2-mercaptoethanol (0.05 mM) at 37°C in 5% CO_2_.

Effector CD4^+^ and CD8+ T cells were generated by stimulating resting CD4^+^ T cells from WT, *Tlr2*^*-/-*^, or *Tlr2*^*fl/fl*^*xCd4*^*cre/cre*^ mice with plate-bound anti-CD3 (1 μg/ml), soluble CD28 (1 μg/ml) with 1 μg/ml of soluble anti-IL4 (Miltenyi) and 10 ng/ml recombinant IL-12 (eBioscience). After three days, cells were washed and rested overnight. On day 4, dead cells and debris were removed by Ficoll gradient centrifugation (Ficoll-Plaque Plus, GE Healthcare), viable cells counted and used for experiments. Resting- or *in vitro*-generated effector CD4^+^ or CD8^+^ T cells (10^5^ cells/well) were stimulated with 1 μg/ml plate-bound anti-mouse CD3 mAb (BD Bioscience) in combination with different concentrations of P3CSK4, LPS (1 μg/ml), anti-CD28 (1 μg/ml), or media alone. Culture supernatants (50 μl) were collected from CD4^+^ and CD8^+^ T cell cultures for IL-2, TNF-α, IFN-γ and IL-10 measurements.

Bone marrow derived macrophages (BMDM) from WT, *Tlr2*^fl/fl^, and *Tlr2*^*fl/fl*^*xCd4*^*cre/cre*^ mice were isolated and differentiated for 7 d in L-929 conditioned media as a source of M-CSF (L-929 cells, ATCC). Prior to use, BMDM were trypsinized, washed, then plated (5×10^4^ cells/well) and allowed to adhere overnight. BMDM were then washed and co-cultured with different concentrations of P3CSK4, LPS (1 μg/ml), or media alone. Culture supernatants (50 μl) were collected from BMDM cultures after 24 h for TNF-α measurements.

### Cytokine quantification

Cytokines were measured by ELISA: IL-2 with anti-IL-2 mAb pairs (eBioscience); TNF-α with BD Bioscience’s TNF-α detection kit (558534); IFN-γ by R&D Systems (SMIF00) and IL-10 by Biolegend (431414)

### Western Blotting

Purified resting splenic CD4+ T cells (2 × 10^6^ cells/mL) were resuspended in cell lysis buffer (20 mM Tris-HCl (pH 7.5), 150 mM NaCl, 1 mM Na_2_EDTA, 1 mM EGTA, 1% Triton), 2.5 mM sodium pyrophosphate, 1 mM β-glycerophosphate, 1 mM Na_3_VO4, 1 μg of leupeptin/mL) and lysed on ice. Cell lysate proteins were resolved on a 4–12% gradient SDS-PAGE gel and transferred to Trans-Blot nitrocellulose membrane (Bio-Rad). Blots were incubated with polyclonal goat anti-mouse TLR2 Ab (R&D Systems; AF1530) followed by HRP-conjugated donkey anti-goat IgG(H+L) mAb (Jackson, West Grove, PA) and bands detected by chemiluminescence using Pierce ECL Plus substrate (Fisher). Blots were analyzed on a Versa Doc Imaging Systems (Bio-Rad) with the Quality One-4.4.0 Software.

### Low-dose M. tuberculosis infection by aerosol

Mice (8-12 wk old) were exposed to aerosolized *Mtb-*H37Rv for 30 min in an inhalation exposure system (Glas Coll, Terre Haute, IN) in the animal BSL-3 facility in order to establish a low dose infection (50-100 CFU/mouse). Animals were sacrificed at different time points, lungs removed with the left lower lobe fixed in 10% formalin for pathology. Remaining lung lobes were placed in Miltenyi C-tubes with 125 U of type IV collagenase and 30 U of DNase/mL. Lungs were gently disrupted using Program Lung 1 of the Miltenyi GentleMacs system, then incubated at 37°C for 30 min. Dissociated lung samples were then further homogenized into a single cell suspension by using Program Lung 2 of the GentleMacs system. Lung homogenates were then clarified with a 40-μm pore-size nylon filter. Red blood cells were lysed and homogenates resuspended in PBS for CFU counting and flow cytometry.

### Flow cytometry

Lung and splenic antigen-presenting cell (APC), CD4+ and CD8+ T cells were analyzed by flow cytometry. Monoclonal antibodies for CD3e, CD4, CD8, CD44, CD62L, CD127, CD8, CD3e, CD25, CD11c f(BD Bioscience), CD4, CD62L, CD44, TLR2 (Ebioscience), I-A/I-E (Biolegend), and Live/Dead Violet (Life Technologies) were used to characterize lung immune cells. Stained samples were acquired using a BD Fortessa flow cytometer. Flow cytometry results were analyzed with FlowJo software (Tree Star, Inc., Ashland, OR). Lung APC were analyzed for MHC-Class II, CD11c and TLR2 expression. Naive, memory, effector and regulatory CD4+ and CD8+ T cells were characterized using CD62L, CD44, CD127 and CD25 expression.

### Statistical analysis

GraphPad Prism version 6 for MacOSX (GraphPad Software, San Diego CA) was used to test for differences in means between specific treatment groups. The null hypothesis of no difference in means between specific treatment groups was tested using a Student’s *t-*test with a p-value <0.05 taken as evidence of a significant difference between groups (\**p* <0.05; \*\**p* <0.01; \*\*\**p* <0.005; \*\*\*\**p* <*0*.*001*).

## Results

### Generation of mice with TLR2 deleted in CD4+ and CD8+ T cells

*Tlr2* floxed C57Bl/6 mice were generated by inserting *loxP* sites to flank *Tlr2* exon 3 (Fig. 1A). Human growth hormone polyadenylation signal (hGH pA) was linked to the 3’ *loxP. Tlr2*^fl/null^ (CWRU0002 TLR2). Heterozygous mice were bred to *Tlr2*^fl/fl^ homozygosity at CWRU. *Tlr2*^fl/fl^ mice were then crossed with *Cd4*^cre/null^ C57Bl/6 mice to generate *Tlr2*-^fl/fl^*xCd4*^cre/null^ C57Bl/6 mice. Littermates of *Tlr2*^*fl/fl*^*xCd4*^*cre/null*^ mice were cross-bred, floxed *Tlr2* verified, and then screened for *Cd4*^*cre*^. To determine *Cd4*^*cre*^ zygosity, mice carrying *Cd4*^*cre*^ were backcrossed to WT mice and litters screened for *Cd4*^*cre*^. If all pups carried *Cd4*^*cre*^, the *Cd4*^*cre*^ carrying parent deemed *Tlr2*^*fl/fl*^*xCd4*^*cre/cre*^. The colony of *Tlr2*^*fl/fl*^*xCd4*^*cre/cre*^ mice was perpetuated, and all experiments were performed with homozygous *Tlr2*^*fl/fl*^*xCd4*^*cre/cre*^ animals. *Tlr2*^*fl/fl*^ and WT mice served as controls.

Thymic precursors of CD4+ and CD8+ T cells express both CD4 and CD8, therefore germline deletion of *Tlr2* in CD4-expressing thymic cells will result in deletion of *Tlr2* in both CD4+ and CD8+ T cells. As shown in Fig. 1B, CD4+ and CD8+ T cells purified from spleens of *Tlr2*^*fl/fl*^*xCd4*^*cre/cre*^ mice contain a DNA fragment that results from deletion of TLR2 exon 3 and is detected by PCR with primers P1 and P3 (Fig. 1A and 1B). Bone marrow-derived macrophages from *Tlr2*^*fl/fl*^*xCd4*^*cre/cre*^ mice have an intact exon 3, resulting in the intervening sequence between P1 and P3 being too large for PCR amplification. CD4+ and CD8+ T cells, and BMDM from WT and *Tlr2*^*fl/fl*^ mice served as additional negative controls (Fig. 1B).

### Impact of Tlr2 deletion on cytokine production by CD4+ and CD8+ T cells

We next determined the impact of deletion of T cell-*Tlr2* on the ability of CD4+ and CD8+ T cells to secrete IL-2, IFN-γ, TNF-α and IL-10, cytokines with a central role in the immune response to *Mtb*. IL-10 has also recently been shown to be highly regulated through TLR2 in non-T reg CD4+ T cells[12]. Resting CD4+ T cells were purified from spleens of WT, *Tlr2*^*fl/fl*^ and *Tlr2*^*fl/fl*^*xCd4*^*cre/cre*^ mice by negative immunomagnetic selection. Purity confirmed by flow cytometry was greater than 95%. Purified T cells were stimulated with plate-bound anti-CD3 mAb in the presence of increasing concentrations of Pam3Cys as co-stimulator. As shown in Fig. 2A, resting CD4+ T cells from WT and *Tlr2*^*fl/fl*^ mice responded as expected to Pam3Cys by increasing IL-2 secretion, whereas CD4+ T cells from *Tlr2*^*fl/fl*^*xCd4*^*cre/cre*^ *mice* did not respond to Pam3Cys co-stimulation. *In vitro*-generated effector CD4+ T cells from *Tlr2*^*fl/fl*^*xCd4*^*cre/cre*^ mice also failed to upregulate IL-2 secretion in response to Pam3Cys co-stimulation (Fig. 2B). In contrast, when stimulated with Pam3Cys, BMDM from *Tlr2*^*fl/fl*^*xCd4*^*cre/cre*^ mice secreted similar levels of TNF-α compared to BMDM from WT and *Tlr2*^*fl/fl*^ mice (Fig. 2C), confirming the selectivity of *Tlr2* deletion in the T cell compartment in *Tlr2*^*fl/fl*^*xCd4*^*cre/cre*^ mice.

**Figure 2.**
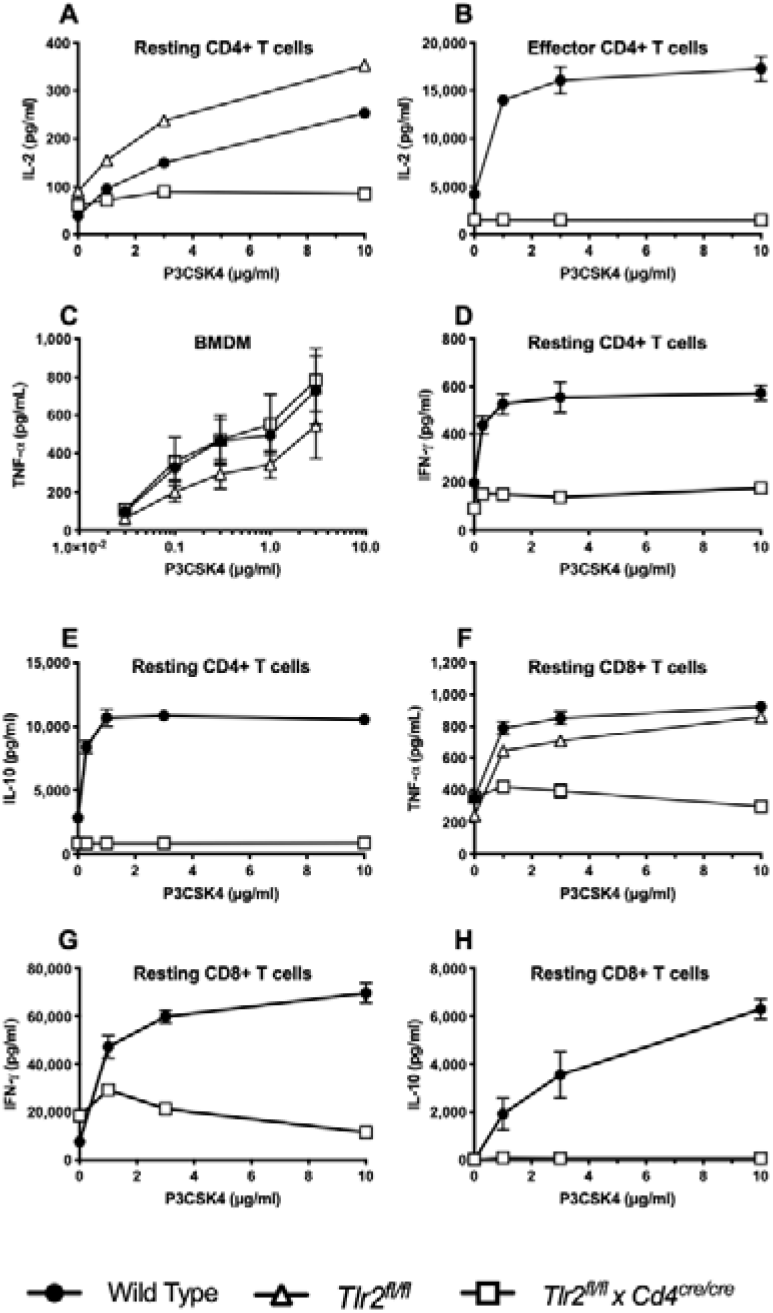
*Cytokine responses to Pam3Cys of CD4+ and CD8+ T cells, and macrophages of Tlr2*^*fl/fl*^*xCd4*^*cre/cre*^ *mice*. Resting and effector CD4+ T cells **(A-D)**, bone marrow derived macrophages (BMDM) **(E)**. and resting CD8+ T cells **(F-H)** were obtained from *Tlr2*^*fl/fl*^*xCd4*^*cre/cre*^, *Tlr2*^*fl/fl*^ and WT mice as described in methods and material. For T cell assays, 10^5^ cells/well in 96 well plates were cultured in triplicate in the presence of plate bound anti-CD3 mAb (1μg/mL) and increasing concentrations of Pam3Cys (0-10μg/mL). BMDM were plated at 5×10^4^ cells/well in 96 well plates in triplicate, adhered overnight, followed by stimulation with Pam3Cys (0-3μg/mL). Supernatants were harvested after 24 h for IL-2 and TNF-α, and 48 h for IFN-γ and IL-10, and measured by ELISA as described in methods and materials. Results presented represent the mean +/-SD of triplicate cultures and are representative of at least 3 experiments per cell type and mouse strain.

We next determined the impact of *Tlr2* deletion on IFN-γ and IL-10 secretion by CD4+ T cells. As shown in Fig. 2D & 2E, Pam3Cys failed to co-stimulate production of these cytokines in CD4+ T cells from *Tlr2*^*fl/fl*^*xCd4*^*cre/cre*^ mice compared to T cells from WT mice. Interestingly, although Pam3Cys co-stimulated cytokine production in CD4+ T cells from *Tlr2*^*fl/fl*^ mice, constitutive IFN-γ and IL-10 production was much higher than that of T cells from WT and *Tlr2*^*fl/fl*^*xCd4*^*cre/cre*^. When checked for T cell TLR2 mRNA and protein expression by RT-PCR and Western blot, no differences were found in TLR2 expression in CD4+ T cells from WT and *Tlr2*^*fl/fl*^ mice (data not shown).

Since little is known about TLR2 as co-stimulator of CD8+ T cells and since *Tlr2* in *Tlr2*^*fl/fl*^*xCd4*^*cre/cre*^ would also be deleted in CD8+ T cells, we determined whether TLR2 functioned as a co-stimulator of CD8+ T cells for cytokine secretion in WT and *Tlr2*^*fl/f*^, and was non-functional in *Tlr2*^*fl/fl*^*xCd4*^*cre/cre*^ mice. As shown in Fig. 2E-H, WT and *Tlr2*^*fl/fl*^ CD8+ T cells are costimulated through TLR2 for TNF-α, IFN-γ and IL-10+ secretion, whereas CD8+ T cells from *Tlr2*^*fl/fl*^*xCd4*^*cre/cre*^ mice fail to upregulate cytokine production in response to Pam3Cys.

### Impact of Tlr2 deletion in CD4+ and CD8+ T cells on Mtb infection in vivo

The next series of experiments sought to determine the impact of *Tlr2* deletion in CD4+ and CD8+ T cells on control of *Mtb* infection in the murine lung. We chose to compare WT and *Tlr2*^*fl/fl*^*xCd4*^*cre/cre*^ mice in light of the above described spontaneous increase in cytokine production noted by T cells from *Tlr2*^*fl/fl*^ mice. By flow cytometry, no differences in the distribution of naïve and memory CD4+ and CD8+ T cells were observed in the lung and spleen at baseline based on CD44, CD62L, CD127 and CD25 expression between WT and *Tlr2*^*fl/fl*^*xCd4*^*cre/cre*^ mice. There were also no differences in MHC-II and TLR2 expression on CD11c+ lung APCs.

However, the CD4+/CD8+ T cells was slightly lower at baseline in *Tlr2*^*fl/fl*^*xCd4*^*cre/cre*^ compared to WT mice in both spleen (data not shown) and lung (Fig. 4C).

**Figure 3.**
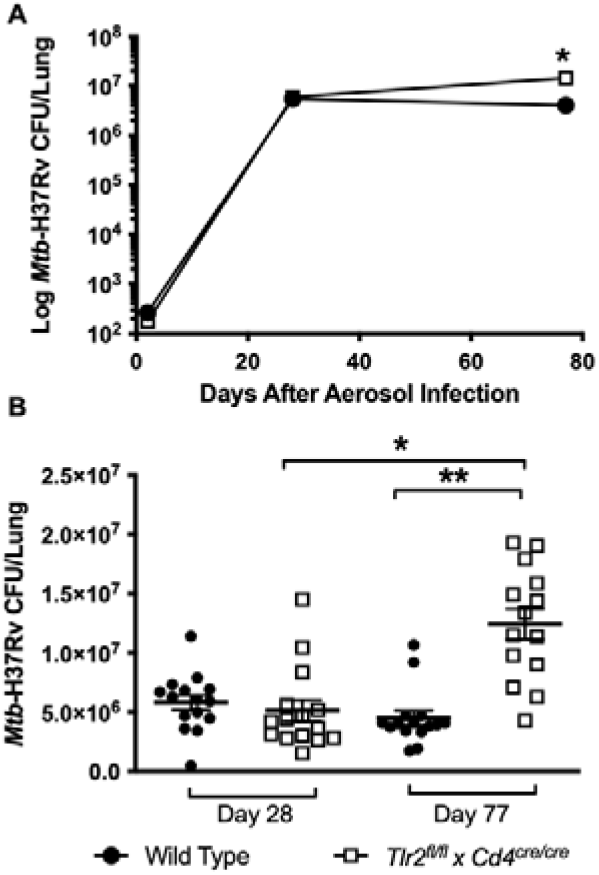
*Mtb growth in lungs of Tlr2*^*fl/fl*^*xCd4*^*cre/cre*^ *and WT mice after low-dose aerosol infection*. Age-matched female mice were infected with *Mtb* H37Rv (50-100 CFU) and lungs harvested 28 and 77 d after aerosol infection. Lungs were homogenized and CFU’s measured as described in methods and materials. **A**. CFUs from a representative experiment involving 5 mice/group, revealing a significant difference (*p <0.05) between the 2 groups of mice. **B**. A summary of CFU from 3 experiments on days 28 and 77, involving 15 WT and 14 *Tlr2*^*fl/fl*^*xCd4*^*cre/cre*^ mice. CFU’s were significantly different (\*\**p* <0.01) between WT and *Tlr2*^*fl/fl*^*xCd4*^*cre/cre*^ at day 77.

**Figure 4.**
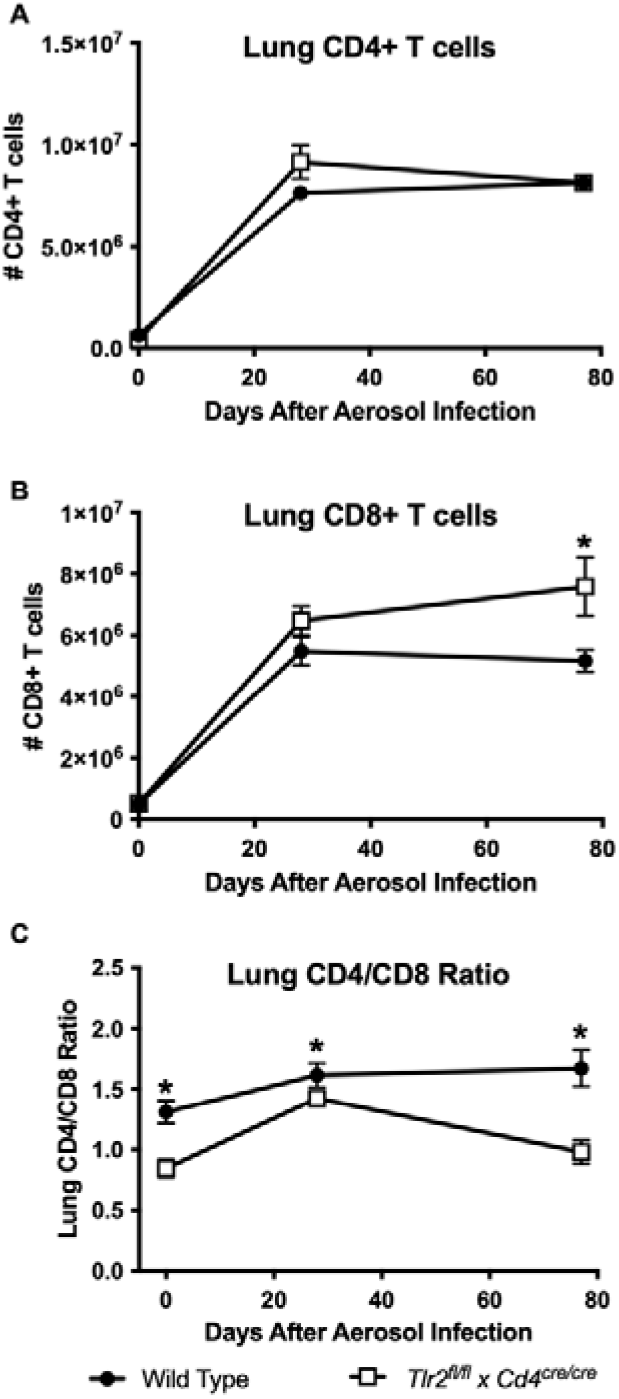
*Distribution of CD4+ and CD8+ T cells in the lungs of WT and Tlr2*^*fl/fl*^*xCd4*^*cre/cre*^ *mice at baseline and during Mtb infection*. Lungs were harvested at different time points, homogenized, cell numbers determined, then stained for CD3, CD4 and CD8 expression before analysis by flow cytometry. Cells were gated on CD3+ events, and CD4+ and CD8+ T cell percentages determined with FlowJo software. **A**. and **B**. Both CD4+ and CD8+ T cell number increased in the lungs of *Mtb* infected mice over time, with CD8+ T cells in *Tlr2*^*fl/fl*^*xCd4*^*cre/cre*^ mice outpacing those of WT mice by day 77 of infection (*p <0.05). **C**. The CD4/CD8 ratio was reduced at baseline in *Tlr2*^*fl/fl*^*xCd4*^*cre/cre*^ mice, and decreased further by day 77 of *Mtb* infection(*p <0.05).

Mice were infected by aerosol with low dose (50-100 CFU/mouse) *M. tuberculosis-* H37Rv. Mice were observed clinically and no obvious differences noted between the 2 experimental groups. Tissues were harvested 28 and 77 d after infection. No differences in CFU/lung were observed after 28 d, but a difference was observed by 77 d with *Tlr2*^*fl/fl*^*xCd4*^*cre/cre*^ mice having a higher *Mtb* burden than WT mice. A summary of CFU results for 3 experiments involving 15 WT and 14 *Tlr2*^*fl/fl*^*xCd4*^*cre/cre*^ mice is shown in Fig. 3A, and a representative experiment is shown in Fig. 3B. The gross appearance of lungs and blinded review of hematoxylin and eosin-stained slides of lung tissue did not reveal differences in lung pathology between the two groups of mice (data not shown). Measurement of lung CD4+ and CD8+ T cell number by flow cytometry showed that the number of lung CD4+ T cells, while increased after infection, did not differ between WT and *Tlr2*^*fl/fl*^*xCd4*^*cre/cre*^ mice at any time point (Fig. 4A).

However, the number of CD8+ T cells in the lungs of *Tlr2*^*fl/fl*^*xCd4*^*cre/cre*^ mice was increased at day 77 compared to WT (Fig. 4B). This increase in CD8+ T cells resulted in a decrease in the CD4/CD8 ratio in the lungs of *Tlr2*^*fl/fl*^*xCd4*^*cre/cre*^ mice by day 77. Thus, selective deletion of *Tlr2* in T cells compromised late control of *Mtb* infection *in vivo*, and this was associated with an increase in CD8+ T cell number and a decrease of the CD4/CD8 ratio in the lung.

## Discussion

The generation of *Tlr2*^*fl/fl*^*xCd4*^*cre/cre*^ mice with selective deletion of *Tlr2* on CD4+ and CD8+ T cells and characterization of their response to *Mtb* described in this manuscript had two goals. First, to determine if *in vivo* activation of TLR2 on CD4+ and CD8+ T cells by *Mtb* ligands impacts control of *Mtb* in the murine lung. Second, since TLR2 signaling impacts macrophage and T cell function, determine where T cell TLR2 contributed to the originally described susceptibility of *Tlr2* gene knockout (ko) mice to aerosolized *Mtb* infection [10, 11].

After generating a floxed *Tlr2* transgenic mouse, a *Tlr2*^*fl/fl*^*xCd4*^*cre/cre*^ mouse was developed resulting in deletion of *Tlr2* in both CD4+ and CD8+ T cells. The baseline distribution of naïve, effector, memory and regulatory CD4+ and CD8+ T cells in lung and spleen were no different between WT, *Tlr2*^*fl/fl*^ and *Tlr2*^*fl/fl*^*xCd4*^*cre/cre*^ mice. There were subtle changes in CD4/CD8 T cell ratios in lungs and spleen of *Tlr2*^*fl/fl*^*xCd4*^*cre/cre*^ compared to WT mice at baseline.

Responses of *Tlr2*^*fl/fl*^*xCd4*^*cre/cre*^ BMDM to TLR-2 ligand Pam3Cys were no different from WT and *Tlr2*^*fl/fl*^ BMDM. We confirmed selective deletion of *Tlr2* on CD4+ T cells by testing with Pam3Cys co-stimulation of cytokine production. We focused on IL-2, IFN-γ TNF-α and IL-10, cytokines of importance to Th1-like immune responses to *Mtb* [2]. We previously demonstrated that mouse and human CD4+ T cells express TLR2 and both synthetic and *Mtb*-derived TLR2 ligands up-regulate IL-2 and IFN-γ secretion by CD4+ T cells [6, 8]. Recent studies have shown that TLR2 co-stimulation also increased IL-10 production by effector/memory CD4+ T cells after 48-72 h [12].

Studies in animal models and humans have established the importance of MHC-I restricted CD8+ T cells in protective immune responses to *Mtb* in the lung [13, 14]. A role for TLR2 on CD8+ T cells in response to *Mtb* is unknown. We found that CD8+ T cells from *Tlr2*^*fl/fl*^*xCd4*^*cre/cre*^ mice lost their ability to upregulate TNF-α, IFN-γ and IL-10 secretion in response to TLR-2 ligand. Recent studies have confirmed a role for TLR2 on CD8+ T cell activation in anti-viral responses and in murine cancer models [9]. Our results demonstrate for the first time that TLR2 also is an effective co-stimulator for CD8+ T cells for cytokine responses relevant to immune control of *Mtb*.

In terms of ability to control *Mtb* growth, we found that deletion of *Tlr2* In CD4+ and CD8+ T cells impacted control of *Mtb* in the lung by 10-11 weeks. These results extend our earlier CD4+ T cell transfer experiments to proving that *in vivo* TLR2 on CD4+ and CD8+ T cells results in enhanced control of *Mtb* [8]. These findings also establish a role for T cell TLR2 in the original studies with TLR2 ko mice demonstrating their enhanced susceptibility to *Mtb* infection. The surprising finding was T cell TLR2’s role in late control of *Mtb*, suggesting that TLR2 costimulation is more important for maintenance of control as opposed to the initiation of *Mtb* control by T cells. The latter is usually apparent by 4-5 weeks. Sugawara *et al*’s finding that TLR2 ko mice were transiently more susceptible after low dose *Mtb* infection after 3 weeks is more consistent with a role for macrophage/APC TLR2 [11]. Reiling *et a*l found that after high dose *Mtb* infection mortality of TLR2 ko mice started to increase by 6 weeks [10]. This finding suggests a role for T cell TLR2 in maintenance but not initiation of *Mtb* control. While TLR2 on macrophages is part of the acute innate inflammatory response to *Mtb*, chronic stimulation of TLR2 inhibits MHC-II antigen processing, allowing *Mtb* infected macrophages to evade CD4+ T cell recognition [5]. The latter is likely relevant during chronic infection since during the acute phase of infection robust CD4+ T cells are generated. The increased late expansion of CD8+ T cells in *Tlr2*^*fl/fl*^*xCd4*^*cre/cre*^ mice in response to *Mtb* infection likely represents compensation for suboptimal CD4+ T cell activation as opposed to contributing to failure of *Mtb* control. Studies in murine models have clearly established the importance of MHC-I restricted CD8+ T cells in protective immune responses to *Mtb* in the murine lung [13, 14].

In summary, our studies demonstrate that deletion of *Tlr2* in CD4+ and CD8+ T cells reduces the ability of these T cells to be co-stimulated for cytokine production by TLR2 ligands expressed by microbial pathogens such as *Mtb*. These include both pro-(IFN-γ, TNF-α) and anti-inflammatory cytokines (IL-10). Furthermore, we found that deletion of TLR2 in T cells *in vivo* did not affect early control but resulted in decreased late control of *Mtb* in the lungs, suggesting that TLR2 co-stimulation of CD4+ and CD8+ T cells *in vivo* is important for maintenance of the steady state between pathogen and host during the chronic phase of *Mtb* infection in the lung.

## Acknowledgements

We thank the CWRU/UH CFAR (NIH P30AI036219) and the Cytometry Facility of the Case Comprehensive Cancer Center (NIH P30 CA43703) for access to core facilities that supported this research. This work was supported by NIH grants AI099494 (to R.E.R), CFAR Developmental Award (CWRU/UH CFAR, P30 AI036219) (to R.E.R), AI034343 (to C.V.H) and AI027243 (to W.H.B.).

